# A novel tamanavirus (*Flaviviridae*) of the European common frog (*Rana temporaria*) encodes a divergent class 1b XRN1-resistant RNA element

**DOI:** 10.1101/2023.07.06.547906

**Authors:** Rhys Parry, Andrii Slonchak, Lewis J. Campbell, Natalee D. Newton, Humberto J. Debat, Robert J. Gifford, Alexander A Khromykh

## Abstract

Flavivirids are small, enveloped, positive-sense RNA viruses from the *Flaviviridae* family with genomes between ∼9-13kb. Metatranscriptomic analyses of metazoan organisms have revealed a diversity of flavivirus-like or flavivirid viral sequences in fish and marine invertebrate groups. To date, however, no flavivirus-like or flavivirid has been identified in amphibians. To remedy this, we investigated the virome of the European common frog (*Rana temporaria*) in the United Kingdom, utilising high-throughput sequencing at six catch locations. De novo assembly revealed a coding-complete virus contig of a novel flavivirid ∼11.2kb in length. The virus encodes a single open reading frame of 3456 amino acids and 5’ and 3’ untranslated regions (UTRs) of 227 and 666nt, respectively. We named this virus Rana tamanavirus (RaTV), as BLASTp analysis of the polyprotein showed the closest relationships to Tamana bat virus (TABV) and Cyclopterus lumpus virus from *Pteronotus parnellii* and *Cyclopterus lumpus*, respectively. Phylogenetic analysis of the RaTV polyprotein compared to *Flavivirus* and Flavivirus-like members indicated that RaTV was sufficiently divergent and basal to the vertebrate Tamanavirus clade. In addition to the Mitcham strain, partial but divergent RaTV, 95.64-97.39% pairwise nucleotide identity, were also obtained from the Poole and Deal samples, indicating that RaTV is widespread in UK frog samples. Bioinformatic analyses of putative secondary structures in the 3′-UTR of RaTV indicated a potential exoribonuclease-resistant RNA (xrRNA) structure identified in flaviviruses and TABV. To examine this biochemically, we conducted an in vitro XRN1 digestion assay showing that RaTV likely forms a divergent but functionally homologous XRN1-resistant xrRNA.

## Introduction

Flavivirids (family *Flaviviridae*) are small enveloped viruses with a positive sense (+), monopartite RNA genome between ∼9-13kb in length and contain the genera *Flavivirus, Pestivirus, Pegivirus*, and *Hepacivirus* (1). Flavivirids have a long evolutionary history and co-divergence with metazoans, with estimates of time scales in the hundreds of millions of years (2-4).

The *Flavivirus* genus can be divided into four ecological groups: the vector-borne vertebrate infecting mosquito (MFV) or tick (TBF) flaviviruses which can cause disease in humans, including the type species yellow fever (YFV) virus (5). Classical insect-specific flaviviruses (cISF) which are restricted to replication in insect cells, and no known arthropod vector flavivirus groups (NKV) (reviewed by Blitvich and Firth (6)). *Flavivirus* genomes encode a polyprotein that is post-translationally cleaved into three structural (capsid (C), premembrane (prM), and envelope (E)) and seven non-structural (NS) proteins (NS1, NS2A, NS2B, NS3, NS4A, 2K, NS4B, and NS5)(7, 8). Additionally, *Flavivirus* genomes contain ∼100 nt 5′ untranslated region UTR and ∼400–700 nt 3′ untranslated regions facilitating viral RNA replication and translation (9). In the flavivirus 3’ UTR, all groups of flaviviruses have three conserved structural RNA elements; the *cis*-acting replication element, a dumbbell, and exoribonuclease-resistant RNA (xrRNA).

Metatranscriptomic and data mining studies reveal a remarkable diversity of viruses infecting eukaryotic species (3, 10). In addition to classical flaviviruses, a wide variety of divergent “flavi-like” or flavivirid viruses or viral sequences have been identified outside terrestrial eukaryotes in several groups of fish (11), marine invertebrates (12) and non-bilaterians (13). Despite sharing conserved protein domain architecture, these flavi-like sequences do not form a monophyletic grouping with the four established groups of flaviviruses, instead representing a basal group to all flaviviruses. These viruses are collectively referred to as tamanaviruses in recent publications (2, 13), named after the prototype species Tamana bat virus (TABV), which was isolated from *Pteronotus parnellii*, an insectivorous bat known as Parnell’s moustached bat (14, 15).

The second vertebrate tamanavirus (VTV), Cyclopterus lumpus virus (CLuV), was discovered in the tissues of a diseased lumpfish (*Cyclopterus lumpus*) (16). Another VTV is Wenzhou shark flavivirus, identified abundantly in all tissues of the Pacific spadenose shark, *Scoliodon macrorhynchos*, from a metatranscriptomic study (3). While genome fragments of similar related viruses have been identified in ray-finned fishes (17, 18), only the Salmon flavivirus from Chinook Salmon (*Oncorhynchus tshawytscha*) has a complete genome sequenced with the full 5’ and 3’ ends of the genome sequenced and been experimentally validated *in vivo* (19).

In addition to tamanavirus genomes, endogenous viral elements or ‘genomic fossils’ of tamanaviruses have been identified in tube-eye fish (*Stylephorus*) and other species (2). This suggests that tamanaviruses - like other groups within the *Flavivirus* family - have ancient origins in the animal kingdom. However, despite extensive studies of the viromes of vertebrates (3), with some targeting amphibians and reptiles (20, 21), a significant phylogenetic gap still exists between aquatic tamanaviruses and vector-borne flaviviruses. This implies that related tamanaviruses may be found in other vertebrates, such as amphibians and reptiles but have yet to be discovered.

We present the first discovery of a tamanavirus, Rana tamanavirus (RaTV), infecting multiple geographically distinct populations of *R. temporaria* frogs in the United Kingdom. Our analysis reveals that RaTV genome organisation is similar to flaviviruses but is genetically divergent and has a dinucleotide composition similar to other vertebrate tamanaviruses. Phylogenetically, RaTV defines a basal position to a putative vertebrate Tamanavirus taxon. We also examined the RaTV genome and identified a genetically divergent XRN1-resistant element (xrRNA/sfRNA) in its 3’ untranslated region and experimentally validated this structure *in vitro*.

## Results

### A novel tamanavirus infects the European common frog (*Rana temporaria*) in UK samples

During the 2014 spring breeding season (February-April) in the UK, toe clip samples were obtained from pairs of male and female adult *R. temporaria* frogs (22) at sites selected from the Frog Mortality Project database and a database of locations known to be free of ranaviral infections (Fig. 1A) (23-26). We conducted high-throughput transcriptome sequencing on these samples, performed *de novo* assembly of the resulting sequences, and screened for virus-like contigs using a previously established pipeline (27).

**Figure 1:**
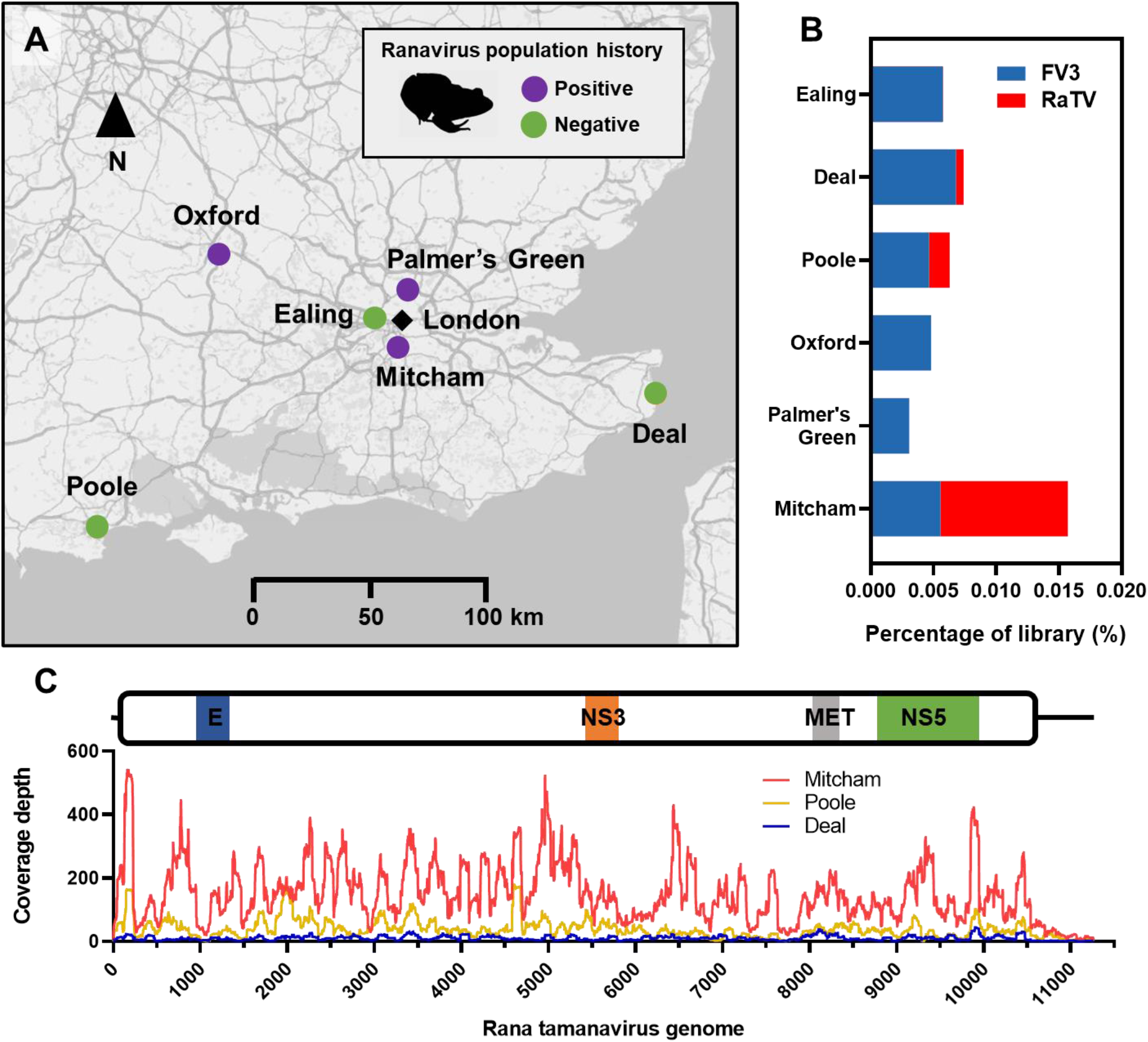
*Rana temporaria* populations sampled for sequencing and incidence of viral infections. A. Map of catch locations in the UK with positive history (purple) and negative history (green) of Ranavirus (FV3) infection. B. Incidence and percentage of sequencing reads for both FV3 (blue) and the novel Rana tamanavirus (red). C. Coverage depth of the RaTV strains from the three catch locations. Flavivirus envelope glycoprotein (E), Non-structural protein 3 and 5 (NS3/5), and methyltransferase (MET) domains are indicated.

Three of the six libraries, Mitcham, Poole and Deal, contained contigs with strong BLASTx hits to the polyprotein of TABV (Accession NP_658908.1, Query cover 86%, Percentage Identity 28.38%, E-value 0, Total score 955) and CuLV (Accession ATY35190.1, Query cover 81%, Percentage Identity 27.94%, E-value 0, Total score 896). The Mitcham library had the largest contig of 11264nt in length. Given the closest hit to this putative virus contig, this virus was named Rana tamanavirus (RaTV). In contrast to CuLV, which encodes a single polyprotein from a predicted programmed −1 ribosomal frameshift (16), the RaTV genome contains a single open reading frame encoding a 3456aa polyprotein and 5’ and 3’ UTRs of 227nt and 666nt, respectively.

We remapped the reads from all libraries to the RaTV Mitcham strain, and also Frog virus 3 (FV3, *Ranavirus*), a dsDNA virus often found in healthy frogs, and found that both disease-free and positive disease history populations contained reads that mapped to the genome of FV3 (Fig. 1B), as previously described (22), while no reads originating from RaTV were found in other libraries. The mapping coverage of the RaTV strains revealed an average coverage of 167x for Mitcham, 57x for Poole, and an average coverage of 10x for Deal. The consensus sequences of the Poole and Deal strains were recovered under stringent coverage and quality scores resulting in 9948nt (88.3%) of the genome recovered for the Poole and 5116 (45.4%) for the Deal strain.

### The Rana tamanavirus polyprotein contains conserved Flavivirus-like protein domains and putative cleavage patterns

In the *Flaviviridae*, the polyprotein is multi-membrane-spanning that becomes embedded in numerous positions in endoplasmic reticulum membranes and is co- and post-translationally processed by the NS2B-NS3 chymotrypsin-like protease(28) and host proteases(29, 30). The resultant conserved proteins include the structural capsid protein (C), pre-membrane protein (prM), and envelope protein (E), as well as nonstructural (NS) proteins NS1, NS2A, NS2B, NS3, NS4A, 2K, NS4B, and NS5.

To annotate the RaTV polyprotein, we predicted the conserved *Flaviviridiae* protein domains, transmembrane topology and protein cleavage motifs (Table 1) in RaTV compared to previous predictions of TABV(15) and CuLV(16) and the *Flavivirus* type species YFV (Fig. 2). RaTV shares conservation of protein domain architecture between putative tamanaviruses and YFV (Fig. 2). Of the structural proteins, RaTV encodes a highly conserved flavivirus E glycoprotein with central and dimerization domains (pfam00869; E-value ≤6.1E-23, Position 304-548). An amino acid alignment of the RaTV glycoprotein with other tamanaviruses and YFV shows that the protein is likely to structurally resemble the domain organisation of prototypical flaviviruses (Supplementary Figure 1). To examine a possible structure of RaTV, the E glycoprotein was modelled using Alphafold2 Colab (Fig 2C). Structures of E in known mature flavivirus virions consist of three ectodomains (DI-III) followed by stem and transmembrane regions anchoring E to the lipid bilayer (31-33). DI forms a central beta-barrel domain which is flanked by DII, an elongated dimerisation domain with the fusion peptide at the distal end. DIII is an immunoglobulin-like domain and is arranged on the opposite side of DI. Each of these domain features were presented in the predicted RaTV glycoprotein structure, including the fusion loop peptide which is highly conserved in RaTV (^382^DRGWTTGCFIFGKGGV^394^) and is essential for viral membrane fusion (34, 35). Regions of low homology between RaTV and typical vertebrate-infecting flaviviruses correlated with disordered predictions in the structure. One example of this is 267-VNVSKHDSFNNEAGSRLAGDYGYSE-291 with a disordered loop protruding from the top of DI, suggesting the structure of E in this region may be substantially different from known flaviviruses.

**Table 1:**
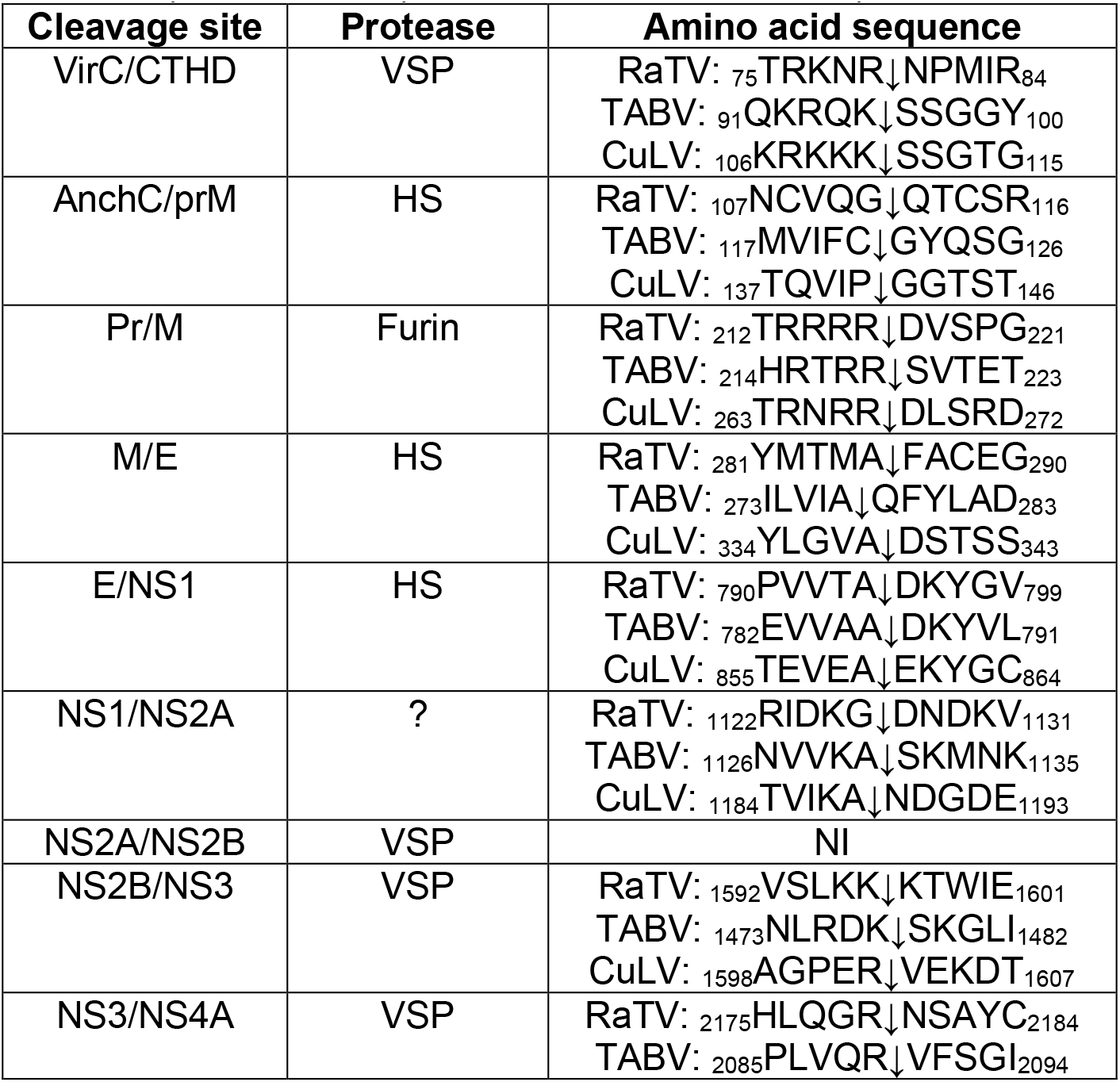
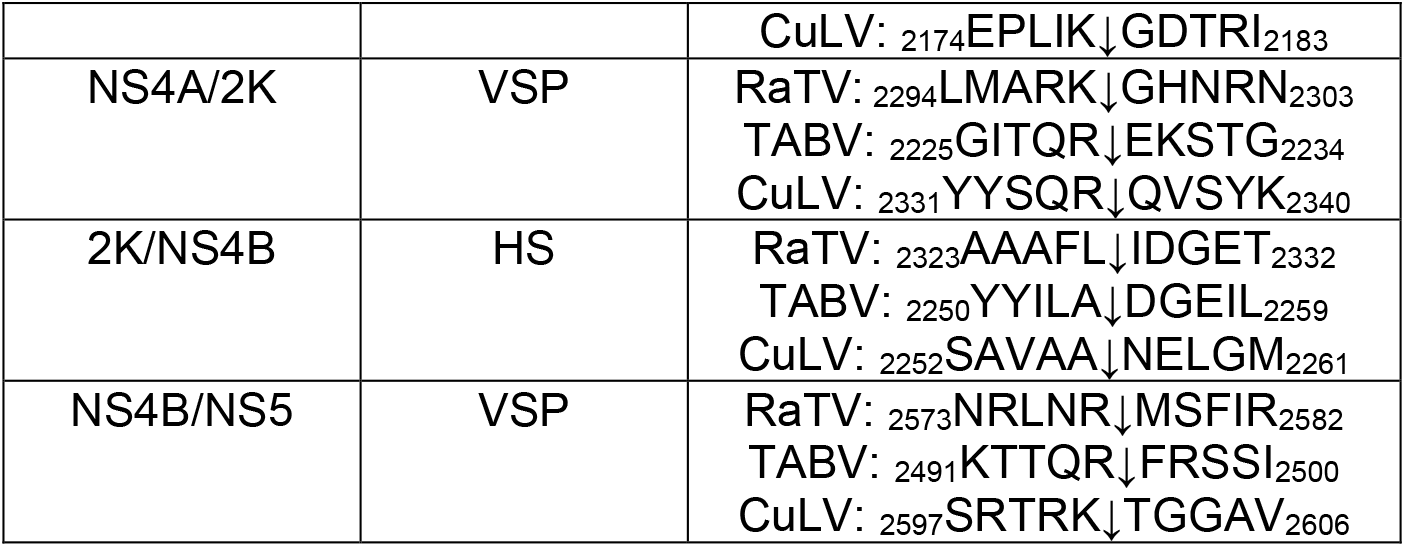
Putative cleavage residues are conserved between vertebrate Tamanaviruses. TABV (GenbankID: NP_658908.1), and CuLV (GenbankID: ATY35190.1)

**Figure 2:**
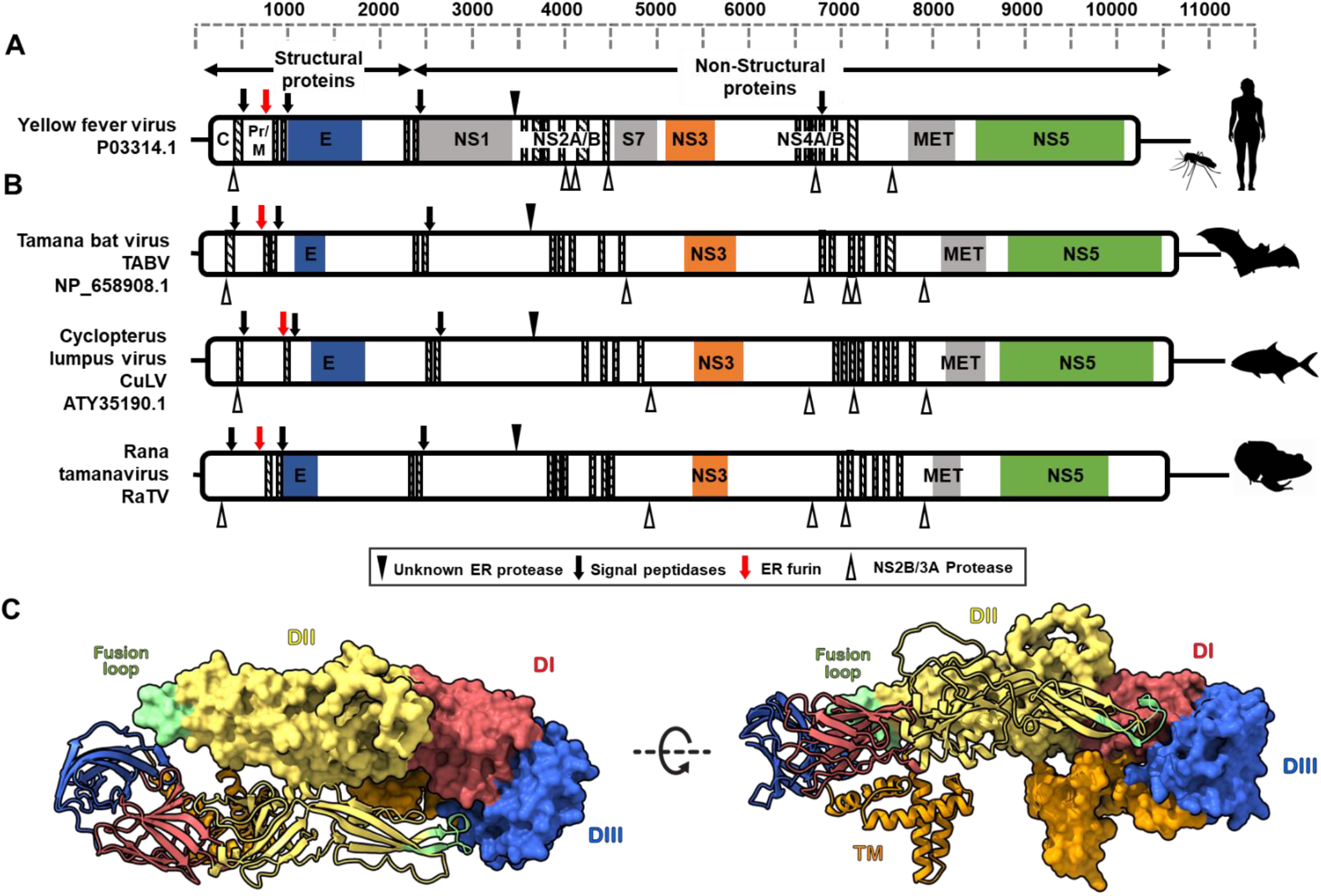
Genome architecture and E protein structure of Rana tamanavirus compared to flavivirus type species. A. Yellow fever virus and B. the putative vertebrate tamanaviruses TABV, CuLV and RaTV. Coloured boxes indicate predicted flavivirus domains: Flavivirus capsid (C) and envelope glycoprotein (E), Non-structural proteins 1-5 (NS1-5), Peptidase (S7) and methyltransferase domain of NS5 (MET). Predicted transmembrane domains are shown with diagonal hatched lines, and predicted polyprotein cleavage residues are indicated as per the legend. C. Cartoon representation of the predicted Alphafold2 RATV E dimer. Residues corresponding to domains are coloured as follows: domain I (DI, red), domain II (DII, yellow) containing a fusion loop (green), domain III (DIII, blue), and E membrane domain containing transmembrane and perimembrane regions (orange).

Non-structural protein domains of RaTV were also highly conserved as RaTV was identified as encoding the conserved flavivirus DEAD domain of the NS3 helicase (pfam07652; E-value ≤7.4E-15, Position 1762-1893) and the helicase conserved C-terminal domain (pfam00271; E-value 0.0073, 1918-2030). The FtsJ-like methyltransferase domain of NS5 (pfam01728; E value 0.026, position 2621-2715) and the flavivirus RNA-dependent RNA polymerase domain of NS5 (pfam00972; E-value ≤6.2E-46, Position 2862-3247). The NS3 serine protease was manually predicted using the conserved trypsin-like serine protease domain catalytic triad (His/Asp/Ser) (Supplementary Figure 2) (36).

Generally, the host furin, putative virus NS2B-NS3-protease cleavage sites and potential host signalase sites for processing RaTV and closely related vertebrate tamanaviruses were conserved. The NS3-protease protein of flaviviruses cleaves after two basic amino acid residues (RR/RK/KR) before a small amino acid (G/A/S).

Although some predicted sites are identical to canonical NS3-Protease cleavage motifs (Table 1), predicting many NS2B-NS3-protease cleavage sites for RaTV and CuLV was challenging as well-established flavivirus residues are weakly conserved in the tamanaviruses (15). Moreover, boxes 3 and 4 of the NS3-Pro domain, which have residues involved in substrate binding and recognition, showed considerable divergence. This may indicate that tamanavirus NS3-Pro may have a different substrate for cleavage that is not the canonical two basic amino acid. The Flavivirus polyproteins pre-membrane (pr) and membrane (M) proteins, processed in the *trans*-Golgi network, are cleaved by the host convertase furin (37), which cleaves at the highly conserved motif R-X-R/K-R. RaTV, TABV and CuLV all have perfect Furin cleavage sites, so it’s likely that the Tamanaviruses process prM in this conserved manner.

### Rana tamanavirus dinucleotide composition is similar to other vertebrate tamanaviruses

Viral RNA containing CpGs are recognised by the vertebrate antiviral protein ZAP, which hinders the multiplication of the virus (38). Classic CpG underrepresentation has been demonstrated in all vertebrate infecting flaviviruses (39). In comparison, there is no clear selection against CpG dinucleotides in classical insect-only flaviviruses (cISF).

The odds ratios, which measure the expected ratio of dinucleotide composition over observed dinucleotide ratios, were calculated to determine the extent of under or overrepresentation of dinucleotide ratios in coding sequences. An odds ratio of ≤0.78 or ≥1.23 indicates statistically significant under or overrepresentation, respectively. The CpG motif in the RaTV virus was found to be significantly underrepresented (0.47) at levels similar to other vertebrate (VTV, n=5) and invertebrate (n=4, ITV) tamanaviruses when compared to insect cISF CpG odds ratios (Fig. 3A).

**Figure 3:**
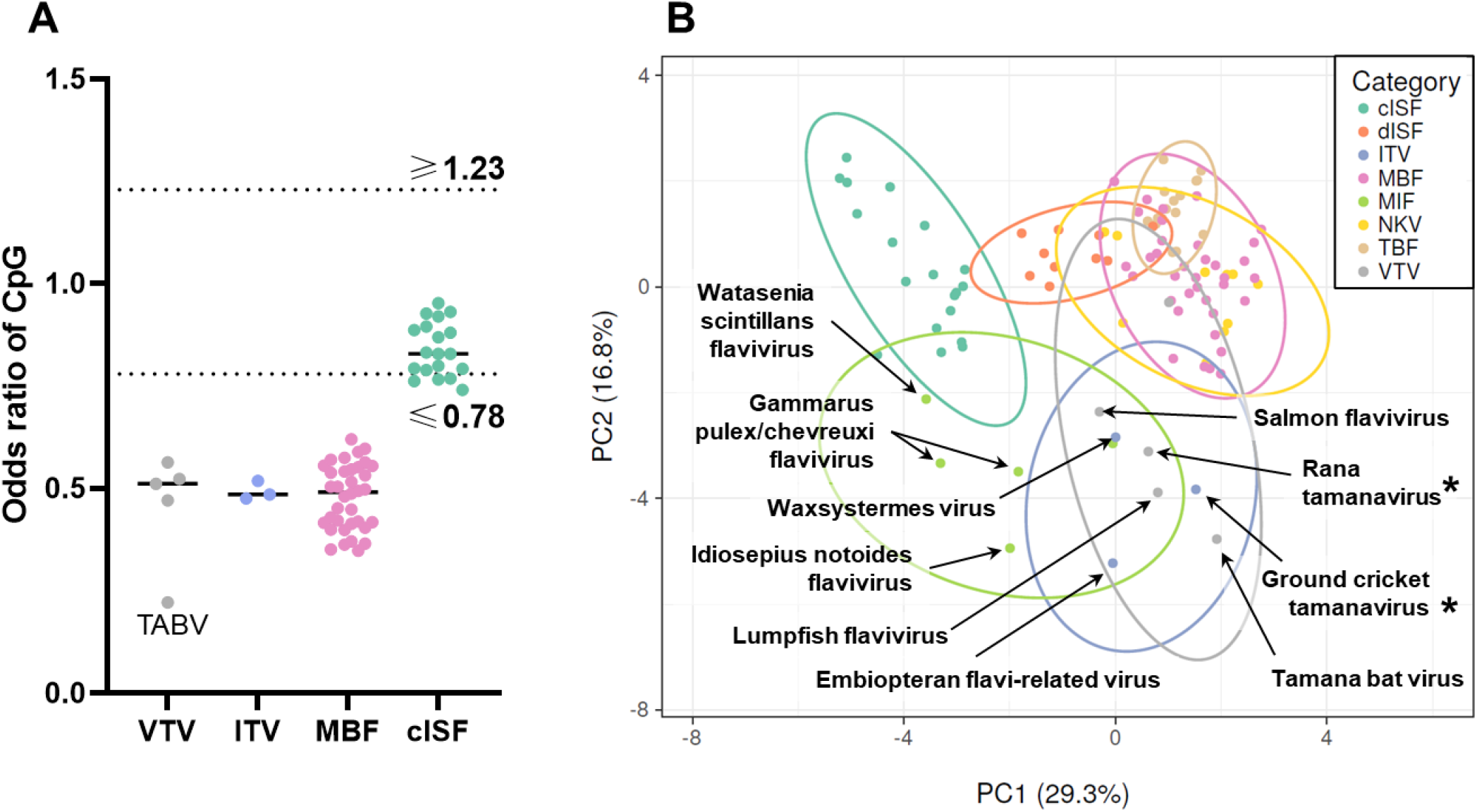
Rana tamanavirus has coding sequence dinucleotide composition similar to other tamanaviruses. A. Odds ratio calculations of the CpG motif of the coding sequence of tamanaviruses and flaviviruses. B. Principal Components analysis of the dinucleotide odds ratios of the coding sequence. Original values are ln(x + 1)-transformed. Unit variance scaling is applied to rows; SVD with imputation is used to calculate principal components. Prediction ellipses bound a probability of 0.95 (n=104). Categories are as follows; cISF: Classical insect-specific Flavivirus, dISF Dual insect-specific Flavivirus, MBF: mosquito-borne flavivirus, NKV: no known vector flavivirus, TBF: tick-borne flavivirus, MIF: marine invertebrate flavivirus, ITV: invertebrate tamanavirus, VTV: vertebrate tamanavirus.

In addition to CpG composition, the collective dinucleotide composition is reasonably predictive of viral hosts in *Flaviviridae* (40). To address the limitations of overinterpreting single dinucleotide motifs, the frequencies of all 16 dinucleotides were used as predictive factors for clustering analyses. Principal component analysis (PCA, Fig. 3B) and hierarchical clustering (Supplementary Figure 3) were conducted to examine the natural groupings proposed for the tamanavirus and *Flavivirus* groups.

The results of the PCA analysis (Fig. 3B) indicate tamanaviruses cluster broadly together but separately from vector borne (MBF, TBF, NKV) and cISF groups. In addition, the dendrogram of hierarchical clustering analysis (Supplementary Figure 3) demonstrates that vertebrate tamanaviruses cluster closer with vector-borne and vertebrate-infecting flaviviruses than the invertebrate tamanaviruses which were in a completely separate group independent of both VTV and MBFs. This indicates that while ITV and VTV may be similar in CpG composition, both groups’ collective dinucleotide composition supports separate host-associated groups independent of the vector-borne flaviviruses and other flavivirus groups.

### Rana tamanavirus is basal to a vertebrate Tamanavirus taxon

To determine the phylogenetic placement of RaTV within the *Flaviviridae* family, 37 representative complete polyprotein sequences were downloaded and aligned from Genbank. This set included several basal flavivirus-like and tamanavirus-like sequences from a variety of insects (41, 42), crustaceans, cephalopods (12), marine vertebrates (3, 16, 19) and TABV (15). We also identified a tamanavirus contig from the transcriptome of the band-legged ground cricket (*Dianemobius nigrofasciatus*, TSA accession: IADE01120511) (43).

Using this alignment, we constructed a consensus maximum-likelihood phylogenetic tree using IQ-TREE2 (Figure 4A). The resultant phylogenetic trees were robustly supported with reasonable bootstrap support values, and this tree’s overall topology is congruent with previously created phylogenies (2, 3). RaTV clusters basally to a clade that encompasses CuLV and TABV. We propose that tamanaviruses can be grouped into two categories based on the host species they infect. The first group, or group one tamanaviruses, includes invertebrate tamanaviruses such as Embiopteran flavi-related, Waxsystermes, and Wenzhou shark flavivirus (3, 12, 41, 42). The second group, or group two tamanaviruses, includes vertebrate tamanaviruses such as RaTV, CuLV, and RaTV.

**Figure 4:**
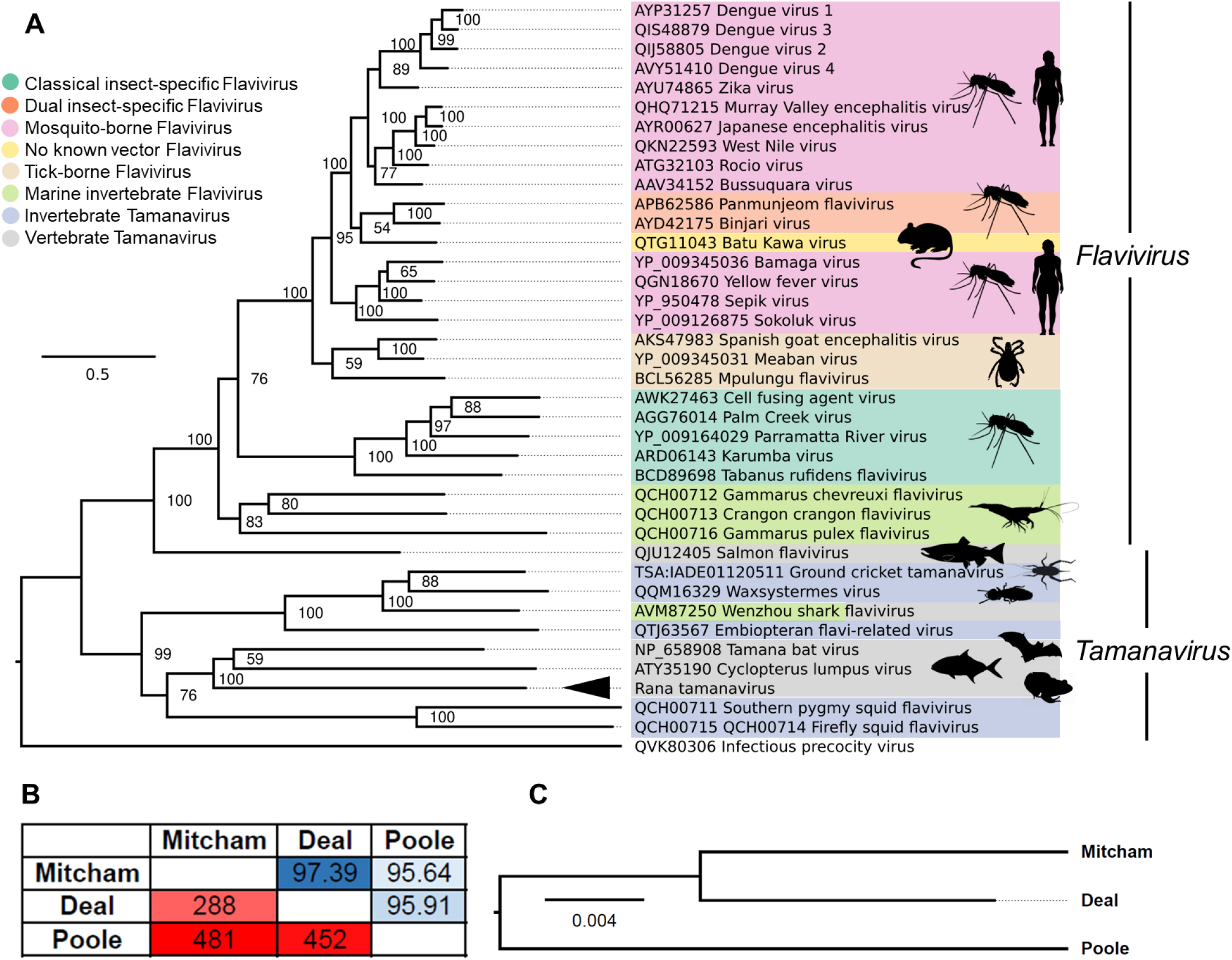
Phylogenetic relationship of Rana tamanavirus within the *Flaviviridae* and Rana tamanavirus strains. A. Maximum clade credibility tree of RaTV compared to multiple ecological *Flavivirus* and the unformalised Tamanavirus taxon. RaTV is indicated with an arrowhead. Branch length represents amino acid substitutions per site with the tree rooted on the outgroup Infectious precocity virus. The GenBank accession of each protein is displayed on the label, and the tree is rooted on the Infectious precocity virus. B. Pairwise nucleotide identity of RaTV strains based on nucleotide differences (bottom left half) and percentage nucleotide identity (top right half). C. Phylogenetic inference of the three assembled RaTV strains based on nucleotide sequence. Branch length represents nucleotide substitutions per site.

To investigate the genetic relationships among RaTV strains from different locations, we assembled and aligned consensus RaTV strains from Deal, Poole and Mitcham retaining 4981nt for analysis. Pairwise comparison of the three RaTV strains revealed a certain level of genetic diversity with 95.64-97.39% nucleotide identity (Fig. 4B). Phylogenetic analysis based on the alignment suggested that the Mitcham and Deal samples were more closely related to each other than to the RaTV strain from Poole (Fig. 4C). These findings indicate that RaTV is prevalent in UK frog samples and over a geographic range of 240km.

### The RaTV 3’ UTR contains a divergent class 1b xrRNA

During flavivirus infection, viral genomic RNA is subjected to degradation by the host 5’-3’ exoribonuclease XRN1 (reviewed by Slonchak and Khromykh (9)). XRN1 is a highly processive enzyme with a helicase activity, which can unwind and fully digest virtually any RNA. However, flaviviruses (44, 45), including TABV (46, 47) contain unique RNA structures called exoribonuclease-resistant RNAs (xrRNAs) within their 3’UTRs that can impede the progression of XRN1. As a result, complete degradation of viral genomic RNA is prevented, and accumulated subgenomic-flaviviral RNAs (sfRNAs) derived from the 3’UTR are present in infected cells. This sfRNA facilitates flavivirus replication by inhibiting antiviral responses in vertebrate and arthropod hosts (48, 49). Production of sfRNA is conserved among all flavivirus clades, while xrRNAs are structurally diverse (45). Three major flavivirus xrRNAs have been described based on their secondary structure (46). Class 1a xrRNAs are found in MBFs, and MBF-like ISFs (lineage 2) (45), class 1b elements are present in classic ISFs (lineage 1), and TABV (45, 46), and class 2 xrRNAs are present in TBFVs and NKVFs (50).

To determine if RaTV contains xrRNAs, we first conducted BLASTn and sequence scan of Rfam analysis of the 3’ UTR. Unfortunately, this analysis indicated no BLASTn similarity with any *Flaviviridae* 3’ UTR or homology with *Flavivirus* Rfam families, including 3′ UTR *cis*-acting replication element (Flavi_CRE; RF00185), *Flavivirus* DB element (Flavivirus_DB; RF00525) or general *Flavivirus* exoribonuclease-resistant RNA element (Flavi_xrRNA; RF03547). Therefore, to identify potential divergent xrRNAs, we then performed a structure and sequence-based alignment of the RaTV 3’UTR with the class 1b xrRNAs of TABV xrRNA (PDB: 7K16_P) (46, 47) and the classical insect-specific flavivirus Palm Creek Virus (PCV) (45) using LocARNA (Figure 5A). This revealed a 48nt element between 11065-11112, approximately 200nt from the terminal end of the RaTV 3’ UTR that displayed homology to known class 1b xrRNA. Subsequent secondary structure modelling (Figure 5B) indicated that this element has the typical organisation of class 1b xrRNAs found in lineage 1 ISFs and TABV, with two coaxially stacked dsRNA helices P1 and P3, a pseudoknot-forming terminal loop L2, side RNA helix P2 with L1 loop, and a small internal pseudoknot. The structure also contained two noncanonical A-C pairs in the base of the P1 helix and was lacking an unpaired C between P2 and P3, which is unique for class 1b xrRNAs. The structure lacked the conserved in 1b class xrRNAs pseudoknot (PK)-forming CAC/G trinucleotide in the apical loop L2. However, putative RaTV xrRNA likely formed a two-nucleotide PK between GC-dinucleotide in L2 and the downstream sequence. This indicates that RaTV and TABV xrRNAs diverged from a common ancestor with cISFs, and clade-specific changes accumulated later in evolution. Given the small and unusual PK in the putative RaTV xrRNA, we then tested the structure for XRN-1 resistance. The *in vitro* transcribed fragment of RaTV 3’ UTR containing the putative xrRNA was treated with the recombinant XRN-1, and the digestion products were analysed by gel electrophoresis. The RNA fragments corresponding to PCV 3’UTR containing known xrRNAs and GFP RNA that is entirely XRN1-susceptible were treated with XRN-1 in parallel to serve as a positive and negative control, respectively. The results demonstrated that XRN-1 could not digest the 3’UTR of RaTV completely and produced a product of incomplete RNA degradation of 200bp (Figure 5C). The size of this product corresponds to the expected size of sfRNA based on the position of the putative xrRNA in RaTV 3’UTR. Similar products of the incomplete RNA digestion by XRN1 were observed with PCV 3’UTR that corresponded in size to the three previously reported sfRNAs of this virus (45). In contrast, the GFP RNA fragment was digested without producing intermediate products. This indicates that RaTV contains a functional xrRNA in its 3’UTR and can produce sfRNA.

**Figure 5.**
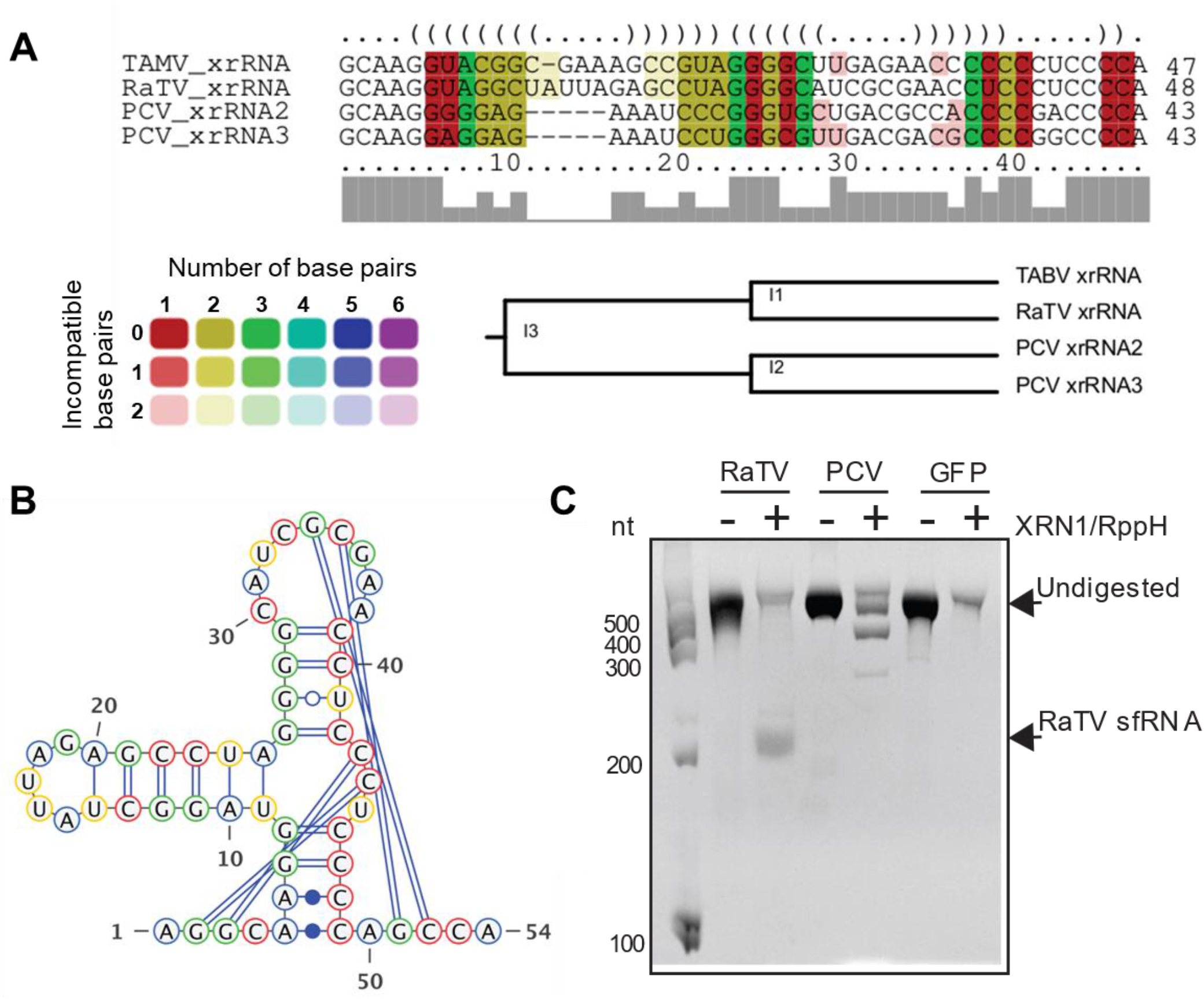
Genetic and biochemical characterisation of the XRN1-resistant RNA element in RaTV 3’UTR. A. Structure-based alignment of RaTV 3’UTR with class 1b xrRNAs of NKVF Tamanavirus (TABV) and cISF PCV performed using LocARNA. B. Predicted secondary structure of RaTV 3’ UTR xrRNA element. Pseudoknots are located manually. C. An *in vitro* XRN1 resistance assay with 3’ UTR of RaTV. RNA fragments corresponding to 3’UTR of cISF Palm Creek Virus (PCV) and GFP ORF are used as the positive and negative controls, respectively. Briefly, RNA was heated, refolded by gradual cooling to 28 °C and then treated with purified XRN1 and RppH (to convert 5’PPP into 5’P). The samples were then analysed by electrophoresis in a denaturing polyacrylamide gel. The gel was stained with ethidium bromide (EtBr).

## Discussion

In this study, we report the discovery of a novel flavivirid/tamanavirus that infects the European common frog. Amphibians are the most threatened group of vertebrates globally, and their populations continue to decline (51) due to a variety of threats, including habitat loss (52), climate change (53), and emerging infectious diseases (EIDs). EID threats to amphibians include fungal chytridiomycosis caused by *Batrachochytrium* (54) and viral pathogens from the dsDNA genus *Ranavirus* (55, 56). Ranavirus infections cause significant adult mortality and the potential for local extinction in *R. temporaria* populations (57), as well as sub-lethal impacts such as altered gene expression profiles (22), changes in skin microbiome structure (58), and shifts in population genetic and demographic structure (57).

The effects of this newly discovered tamanavirus on its host and its mode of transmission are currently unknown. Although, RaTV infection is likely systemic as the virus was detected from toe clips. It is worth noting that for the related TABV, the virus has been readily recovered from the salivary glands and lungs of bats with naturally acquired infections (59). Experimental infections of closely related bat species resulted in productive infections that were isolated from sera, spleens, salivary glands, and saliva (59), with antibodies to TABV detected by hemagglutinin inhibition and neutralization tests at six days. While signs of illness were not observed in any bat, experimental infections of a white-faced capuchin monkey (*Cebus nigrovittatus*) revealed behavioural signs of illness. Infection of infant mice with TABV by the intracranial and intraperitoneal routes was also fatal. While RaTV may be a latent or asymptomatic infection in *R. temporaria* frogs, given the potential for coinfection of RaTV with ranaviruses and other EIDs, the complex interactions between these viruses should be examined.

In addition to showing that RaTV has divergent but structurally conserved flavivirus-like protein domains, we can demonstrate that RaTV encodes an xrRNA-resistant element in its 3’ UTR. Two copies of xrRNA elements exist in the 3’ UTR of TABV (Genbank Accession: MZ229974), with the tertiary structure recently solved by X-ray crystallography (47). TABV xrRNA shares conserved features from previously characterized xrRNAs and also a novel set of tertiary interactions (47), informing later searches for undiscovered xrRNAs. The discovery that the same structural RNA can be achieved by very different sequences and interactions in both the Tamanaviruses and distantly related flaviviruses have provided insight into the convergent evolution and diversity of structural RNAs within this family. While only the xrRNA elements have been characterised, additional 3’ UTR elements, such as potential *Flavivirus* DB elements and terminal 3’ stem-loops, should be examined as additional vertebrate Tamanaviruses also get discovered.

In our recent study, we demonstrated that xrRNAs of this class are present in 3’UTRs of all analysed classical insect-specific flaviviruses (cISFs) (45). Although xrRNAs of this type are common in ISFs, they are relatively rare in other flavivirus clades. For example, they don’t occur in MBFs or TBFs, and within the NKV clade, they have been only found in TABV, while other NKVFs and TBFs analysed to date contained even more divergent class 2 xrRNAs. Herein we found that RaTV contains xrRNA homologous to that of TBAV and cISFs, but divergent from xrRNAs of other characterised NKVs.

In the absence of an authentic viral isolate, alternative methods can be employed to recover RaTV *in vitro* for downstream characterisation. Given the recovery of large and potentially complete portions of the 5 and 3’ UTR, one possible approach utilising reverse genetics could be attempted using synthetic gene blocks tiling the RaTV genome (60) and *de novo* generation with circular polymerase extension reaction (CPER) (61). The resulting CPER product could be transfected into a suitable, highly susceptible mammalian cell line such as VeroE6, African green monkey kidney cells used to propagate TABV. If it is not possible to recover a full-length infectious virus, the structural and antigenic properties of RaTV could be assessed by incorporating the prM-E sequence into the Binjari virus (BinJV) recombinant virus platform (62). The BinJV platform is a lineage two insect-specific flavivirus, which is mosquito-restricted but allows for the generation of antigenically authentic virion particles (62).

## Methods

### Ethics statement

This project was approved and conducted in compliance with ethical guidelines set by the Zoological Society of London and the University of Exeter. Field sampling was conducted under Home Office license 30/10730, issued to Lewis Campbell, and a project licence 80/2466.

### Field site selection and sample collection

During the 2014 spring breeding season (February–April) in the UK, pairs of adult *R. temporaria* frogs were captured during mating, and tissue samples were collected within 24 hours. Before sampling, frogs were rinsed in sterile water, and a disinfectant with analgesic (Bactine; WellSpring Pharmaceutical, Florida, USA) was applied to the distal portion of their hind limbs. Tissue samples were collected by licensed personnel who, after removing frog toe clips, placed the samples in RNAlater (Sigma Aldrich, Missouri, USA). The number of frogs sampled at each site ranged from 2 to 18. After sampling all animals were released.

#### RNA extraction, sample pooling and RNA sequencing

Toe clips from sampled *R. temporaria* frogs were homogenized and lysed using a Qiagen High-Frequency Tissue Lyser2 (Qiagen, Hilden, Germany) with lysis buffer and stainless-steel lysis beads at 2,000 Hz for 4 min. RNA was extracted from the samples using RNA isolation kits from Macherey-Nagel (Duren, Germany) following the manufacturer’s instructions. Extracted RNA was quantified using NanoDrop and evaluated for quality using a BioAnalyser (Agilent Technologies, California, USA). Only samples with RNA integrity scores greater than eight were chosen for pooling. Six individuals (three males and three females) from each site were pooled. The RNA concentrations of each individual within a given pool were normalized to the lowest individual, but the resulting six pools were not normalized between each other. Reverse transcription was conducted using a Superscript II kit (Invitrogen, Massachusetts, USA), a combination of random hexamer and oligo DT primers. Libraries were sequenced on a HiSeq2000 using v3 chemistry, generating 100 base pair paired-end reads.

### *De novo* assembly and phylogenetic analysis of Rana tamanavirus

Adapter sequences and low-quality reads were removed from individual fastq libraries from the six site-specific pools using Cutadapt v1.21(63). Clean reads were assembled using rnaviralSPAdes (v3.15.4) (64). The SPAdes assembled contigs were queried against a previously constructed non-redundant virus protein database(27) using BLASTx (65). The polyprotein of the RaTV Mitcham strain was predicted using the Open Reading Frame Finder (https://www.ncbi.nlm.nih.gov/orffinder/).

For phylogenetic placement within *Flaviviridae*, 37 representative polyprotein sequences were downloaded from Genbank, and one identified Tamana-like viral sequence from the Transcriptome Sequence Archive (Accession: IADE01120511 Ground cricket tamanavirus) and the outgroup Infectious precocity virus strain ZJJS2019 (66) were aligned using MAFFT (v7.490, FFT-NS-I method, BLOSUM45) (67). Ambiguous blocks were subsequently removed from the alignment using Gblocks (v0.91) (68), resulting in a 39 × 1748aa multiple sequence alignment. IQ-TREE2 (v2.1.2) was used for protein substitution model testing and constructing a consensus maximum-likelihood phylogenetic tree with ultrafast bootstrap and SH-aLRT test (--alrt 1000 -B 1000). The LG+F+R6 protein substitution model was selected for alignment using the Bayesian Information Criterion in ModelFinder (69). Phylogenetic tree files were visualised using Figtree v1.4.4 (A. Rambaut; https://github.com/rambaut/figtree/releases).

To recover the Poole and Deal strains of RaTV from the sequencing data, clean fastq base-called files were mapped to the Mitcham strain using Bowtie2 (v2.2.7) under default settings (70). The depth of coverage of mapped alignment files was determined using samtools (v1.3) depth as described previously (71). Single-nucleotide variants of alignment files were identified using iVar (v1.2.2) (72) with a minimum quality score threshold of 20. Consensus positions were called with a minimum depth of 6. After consensus sequences were generated, pairwise comparisons between nucleotide identities were created using CLC Main Workbench (v6.9.2). A maximum-likelihood phylogeny between the three strains was generated using IQ-TREE2 using the model GTR+I+R and previously described parameters. The number of reads originating from each sample for FV3 was also determined using Bowtie2 as above and the reference accession OM927978.1.

### Rana tamanavirus functional genomic annotation and E protein structural modelling

To identify the functional domains in the predicted RaTV polyprotein, we conducted a protein domain-based search using InterProScan (v5.60-92.0, available at https://www.ebi.ac.uk/interpro/search/sequence/). To predict cleavage residues of the polyprotein, we used a transmembrane topology prediction using the TMHMM Server v. 2.0 (https://www.cbs.dtu.dk/services/TMHMM/). To identify putative signal peptides, a 40-60aa sliding window of the polyprotein was assessed by the SignalP v6 webserver (https://services.healthtech.dtu.dk/service.php?SignalP-6.0). To analyse dinucleotide motifs of the coding region of the RaTV genome, we added to a previous dataset of the odds ratio of dinucleotide motifs 103 *Flaviviridae* genomes deposited in Genbank (12). Hierarchical clustering and principal components analysis was performed using the ClustVis(73) web server (https://biit.cs.ut.ee/clustvis/). The original values were ln(x + 1)-transformed before the analysis.

The RaTV envelope glycoprotein monomer was modelled by ColabFold v1.5.2 (74), which utilises AlphaFold 2 (75) using the templates in pdb70 and MMseqs2 under default conditions. The five top models were ranked based on a per-residue confidence score (pLDDT) (Supplementary Figure 4) and manually inspected using ChimeraX (v1.5) (76). Model number 1 was chosen for downstream visualisation. The dimer was arranged using PDB:7KV8, with the domains of E coloured according to amino acid alignments and arrangement in the putative structure.

### XRN1 resistance assay

A DNA fragment corresponding to RaTV 3’UTR under the T7 promoter was synthesised and cloned into the pIDT vector (IDT, Singapore). Plasmids containing PCV 3’UTR and GFP gene fragments have been described previously (45). For *in vitro* transcription plasmids were first linearised by restriction digest and purified using Monarch PCR and DNA Clean-up Kit (NEB, USA). 3’UTRs were *in vitro* transcribed from 500ng of plasmid DNA using MEGAscript T7 Transcription Kit (Invitrogen, USA). RNA was purified by LiCl precipitation and examined by electrophoresis in a 1% denaturing agarose gel. RNA was then refolded in NEB3 buffer (85 °C for 5 min) and gradually cooled to 28 °C. For XRN1 treatment refolded RNA (1 μg) was incubated with 1U XRN1 (NEB, USA) and 10U RppH (NEB, USA) in 20 μL of reaction mixture containing 1x NEB3 buffer (NEB, USA) and 1 U/μL RNasin Plus RNase Inhibitor (Promega, USA) for 2 h at 28 °C. The reaction was stopped by adding 20 μL of Loading Buffer II (Ambion, USA), heating for 5 min at 85 °C and placed on ice. The denatured RNA samples were loaded into 6% polyacrylamide TBE-Urea gels (Invitrogen, USA), and electrophoresis was performed for 90 min in 1xTBE. The gels were stained with ethidium bromide and imaged using an OmniDoc imager (Cleaver Scientific, UK).

### RNA structure prediction

The secondary structures for the RaTV 3’UTR elements were predicted using mfold v2.3 (http://www.unafold.org/mfold/applications/rna-folding-form-v2.php) (77) with parameters: folding temperature 28 °C and maximum distance between paired nucleotides limited to 150. The secondary structures were visualised using VARNA v3.93, noncanonical C-A pairing was manually forced, and pseudoknots were located manually. Structure-based multiple alignments of xrRNAs were performed using RIBOSUM-like similarity scoring implemented in the LocARNA package (v 1.9.1)(78). Experimentally determined (PCV and TABV) or predicted secondary xrRNA structures (RaTV) were provided as structural constraints (#S option).

## Data availability statement

Raw high throughput sequencing read files are archived at the NCBI Sequencing Read Archive database under BioProject ID PRJNA431814 and accession numbers SRR6515974-SRR6515979. The Rana tamanavirus Mitcham strain sequence is available in Genbank under the accession OQ164658. The RaTV Alphafold E protein model is available from Figshare doi: 10.6084/m9.figshare.22179512.

## Acknowledgements

This work was funded by the Australian Research Council (ARC) grant DP190103304 to AAK and AS. and National Health and Medical Research Council (NHMRC) Ideas grant 2021272 to AS. NDN contributions to the work were funded by NHMRC Investigator grant APP2009707. This work utilized compute resources from the Galaxy Australia server (https://usegalaxy.org.au/). L.J.C’s contributions to the work were funded by two sources: a PhD stipend from the Natural Environment Research Council and an Intra-European Fellowship from the FP7 People: Marie-Curie Actions, both awarded and supervised by Dr. AGF Griffiths, University of Exeter. We want to acknowledge Trenton W. J. Garner and Amber G. F. Griffiths for their contributions to original data generation, and the Garden Wildlife Health project.

---

**Supplementary Figure 1.**
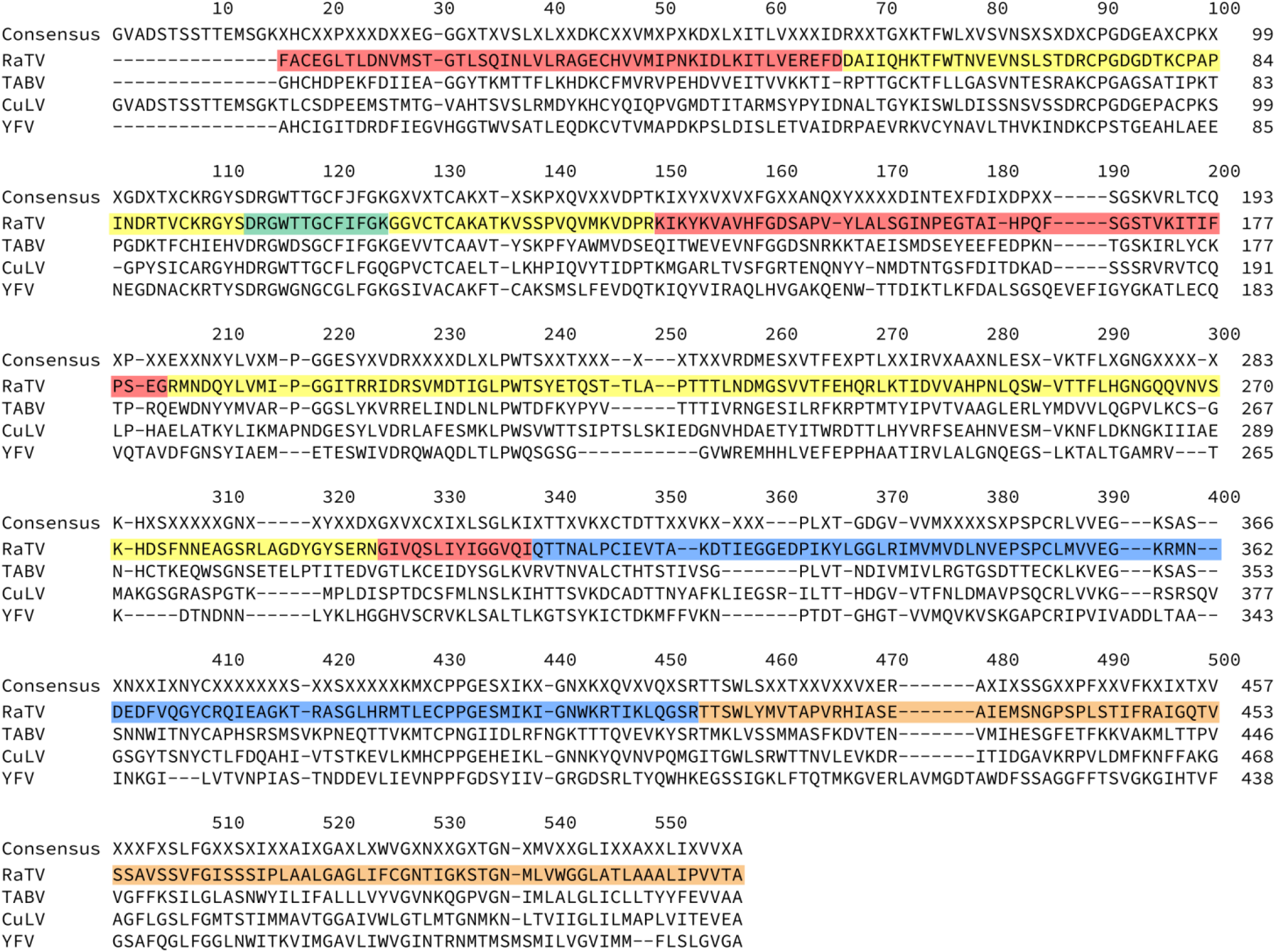
Alignment of the flavivirus E glycoprotein with the prototype *Flavivirus* species YFV (GenbankID: P03314.1), TABV (GenbankID: NP_658908.1), and CuLV (GenbankID: ATY35190.1). Residues are labelled as per domains identified in Figure 2; with the domain I (red), domain II (yellow), domain III (blue), fusion loop (green) and transmembrane and stem region (orange).

**Supplementary Figure 2.**
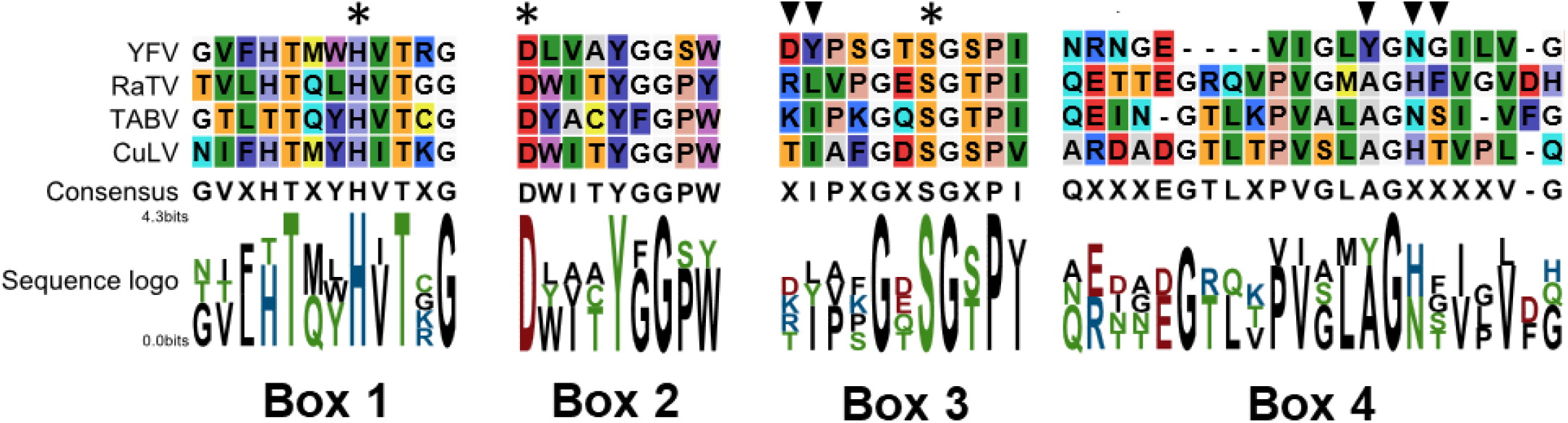
Alignment of the four conserved trypsin-like serine motif boxes of the NS3-Pro region of Rana tamanavirus. The prototype Flavivirus species YFV (GenbankID: P03314.1), TABV (GenbankID: NP_658908.1), and CuLV (GenbankID: ATY35190.1). Arrowheads indicate putative substrate-binding sites, and trypsin-like serine protease catalytic triad residues are indicated with an asterisk (*).

**Supplementary figure 3.**
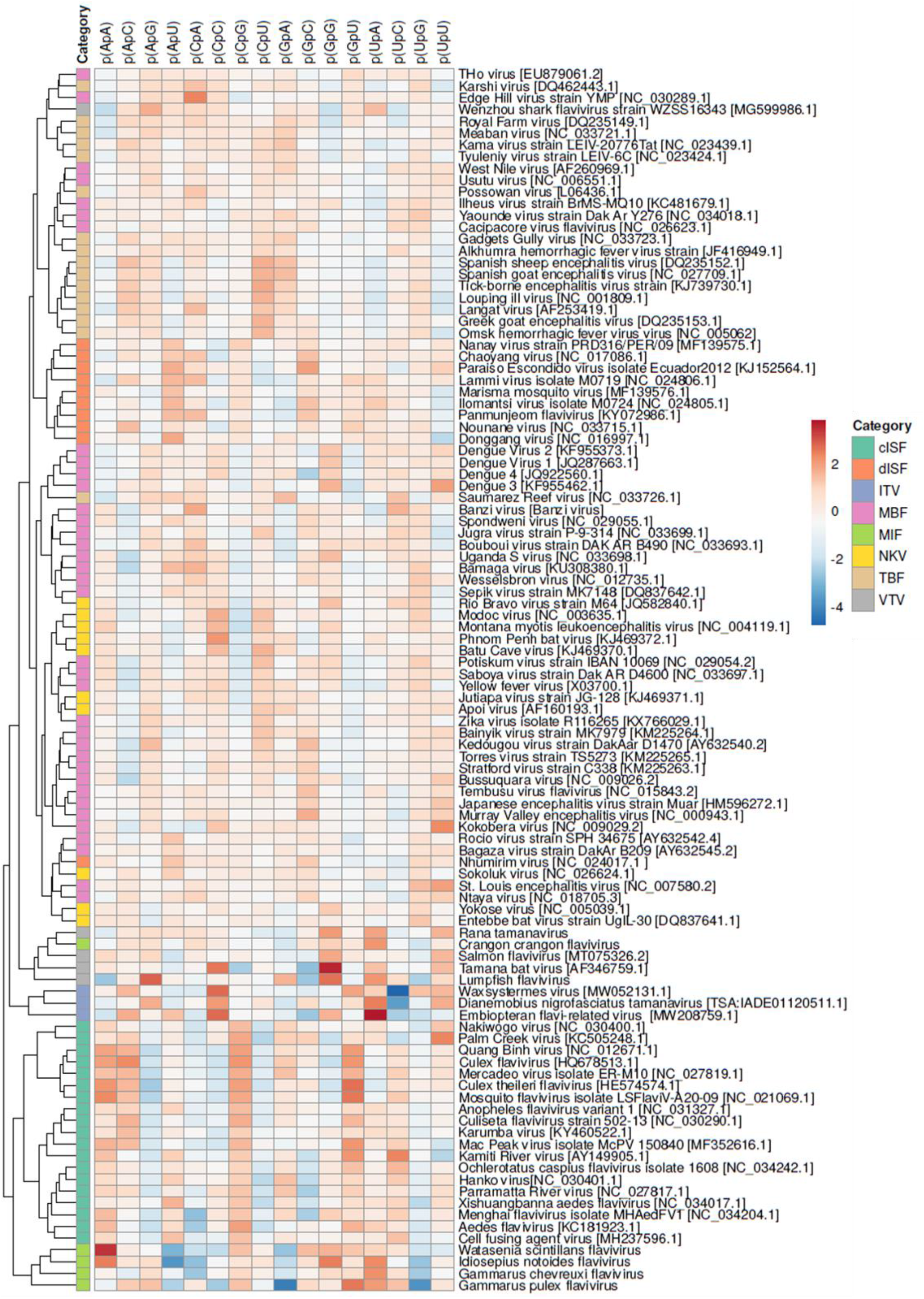
Hierarchical clustering analysis of dinucleotide composition of *Flavivirus* and flavivirus-like tamanaviruses. Rows are centred, and unit variance scaling is applied to rows. Columns are clustered using Euclidean distance and complete linkage. Original values are ln(x + 1)-transformed. Categories are as follows; cISF: Classical insect-specific Flavivirus, dISF Dual insect-specific Flavivirus, MBF: mosquito-borne flavivirus, NKV: no known vector flavivirus, TBF: tick-borne flavivirus, MIF: marine invertebrate flavivirus, ITV: invertebrate tamanavirus, VTV: vertebrate tamanavirus.

**Supplementary Figure 4.**
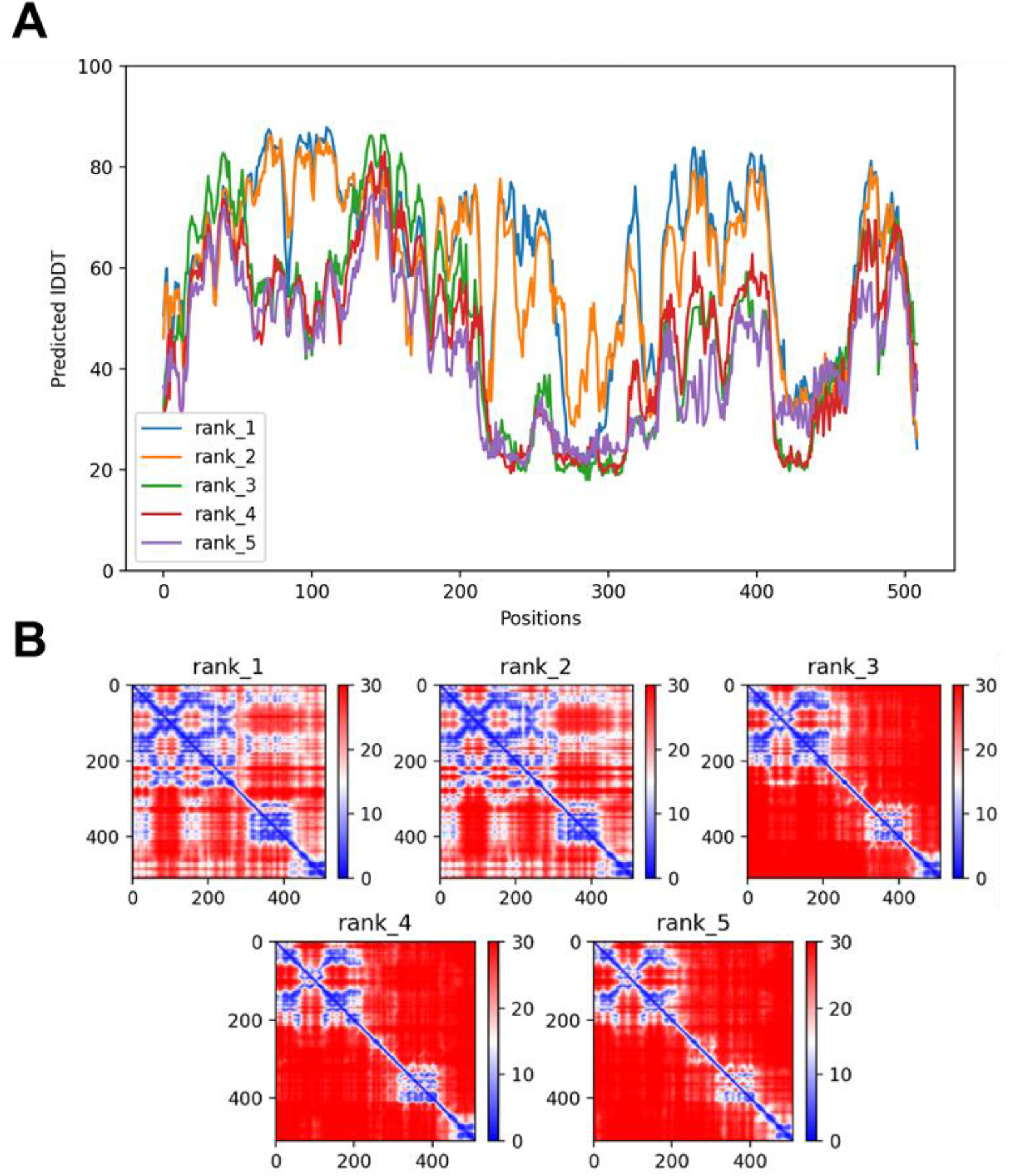
Confidence metrics and quality assurance of AlphaFold structure predictions for RaTV E protein. (A) Local distance difference test (LDDT) values for the five candidate predicted models of RaTV. LDDT is a superposition-free score that evaluates local distance differences in a model compared to a reference. B) Predicted alignment error (PAE) plot, showing the predicted deviation from modelled positions for each residue pair in the AlphaFold model shown for the top five ranked models.

